# Improving long-read somatic structural variant calling with pangenome and de novo personal genome assembly

**DOI:** 10.1101/2025.10.28.685154

**Authors:** Qian Qin, Jakob Heinz, Heng Li

## Abstract

Accurate detection of mosaic and somatic structural variants (SVs) provides early diagnostic and therapeutic evidence for cancers. While long-read whole-genome sequencing leads to more accurate SV detection than short read sequencing, existing long-read SV callers only look at alignment against a single reference genome and are susceptible to systematic false discovery caused by germline differences between the individual genome and the reference genome. Here we develop a new SV calling method that jointly considers the alignment against a pangenome and the de novo assembly of the germline genome. It dramatically reduces false positive mosaic SVs in normal samples and somatic SVs in cancer cell lines with little loss in sensitivity. Our study highlights the essential need for pangenome or personal genome assembly to integrate SV calls for both SV discoveries and clinical diagnostics.

**Significance:** We introduced a novel long-read SV calling method that leverages pangenome and personal genome and greatly improves the accuracy of somatic SV calling.

## Introduction

Structural variants (SVs) are a broad spectrum of genetic variants longer than 50bp, including inter-chromosomal translocations. They can be classified as simple types such as deletion, insertion, inversion, duplication, and translocation, or complex types involving multiple variations in one mutation event such as chromoplexy and chromothripsis(1). Replication stress, virus integration and cellular processes to repair double strand DNA breaks such as homology directed repair can mechanistically lead to most of the SV events(2). These SV events form major sources of genome mutation and instability hallmark in cancer(3). A recent pan-cancer Whole Genome Sequencing (WGS) project has enabled a more comprehensive overview of the somatic SVs prevalence in cancer and discovered that 55% of driver mutations represent SVs, and driver SVs are more prevalent than point driver mutations in some cancers such as breast and ovary adenocarcinomas(2,4,5).

Traditional techniques to identify SVs, such as FISH (fluorescence in situ hybridization), have low-throughput and can only analyze a few loci. Short-read WGS has genome-wide coverage and can capture SVs across the genome. However, it has notable genome coverage bias(6,7) and alignment uncertainty for repeat-rich or highly homologous regions near the telomere, centromere and the variable number of tandem repeats (VNTR) area. Long-read sequencing, as nominated as the year of method in 2022(8), is promising to fill these gaps with reduced coverage bias(9) and improved mappability.

Several long-read SV callers have been optimized for somatic SV calling for tumor-normal pairs, including Severus(10), SAVANA(11), SVision(12), Sniffles2(13) and NanomonSV(14). They take a tumor read alignment and a normal read alignment as input to find SVs specific to the tumor sample. All these SV callers only consider alignments against a single reference genome. At a germline insertion/deletion (indel) relative to the reference, it is possible that a small number of tumor reads are misaligned due to the indel and look like a different SV, but normal reads are not misaligned by chance. This would lead to a potential false somatic SV call as somatic events are generally supported by fewer reads. Current SV callers have to introduce heuristics, such as filtering out VNTR regions, to alleviate the issue, but these hacks are not solving the root problem: false somatic SV calls caused by misalignment around germline SVs.

Mosaic SV calling from normal tissues is even more challenging. It is hard to distinguish mosaic and germline SVs in the lack of a sample without mosaic events. At present, only Sniffles2 calls mosaic SVs. However, it calls hundreds of mosaic SV calls in normal samples, the great majority of which are false positives. There are no capable somatic or mosaic SV callers without matched normal samples.

In this article, we will describe a new method to greatly reduce false somatic or mosaic SV calls caused by germline SVs. This method jointly considers alignment against the de novo assembly of the normal sample or against a pangenome, in addition to the alignment against the single reference genome. Because an ideal de novo assembly encodes all germline SVs and a pangenome represents most common germline SVs, we can reduce misalignment caused by germline SVs and thus improve somatic or mosaic SV calling accuracy. Our method can be applied independently or used jointly with existing SV callers. Evaluated on five normal human samples as true negative cases and six tumor-normal pairs of cell lines as true positive cases, the method greatly reduces false positive somatic SV calls at little cost on sensitivity.

## Materials and Methods

### Method overview

Minisv is a command-line tool that calls somatic SVs, filters existing SVs, evaluates SV calls and constructs the union call set from multiple callers. Minisv is unique in that it can jointly consider the read alignment against multiple genomes or pangenomes to reduce alignment artifacts caused by germline SVs not present in a single reference genome. This feature is particularly powerful when the normal sample in a matched tumor-normal pair can be de novo assembled.

As a standalone SV caller, minisv identifies SVs in three steps (Fig. 1): 1) for each read, extract candidate SVs from its alignment against the linear reference genome, a pangenome and the self-assembly of the normal sample if available; 2) for each read, compare candidate SVs extracted from each alignment and only retain an SV if it is supported in all alignments; 3) merge remaining candidate SVs across reads based on their types and sizes and their positions on the reference genome. This procedure leads to a non-redundant SV callset. For other SV callers that output read names, minisv can use a variant of step 2) to filter their somatic SV calls not supported by the normal self-assembly.

**Figure 1.**
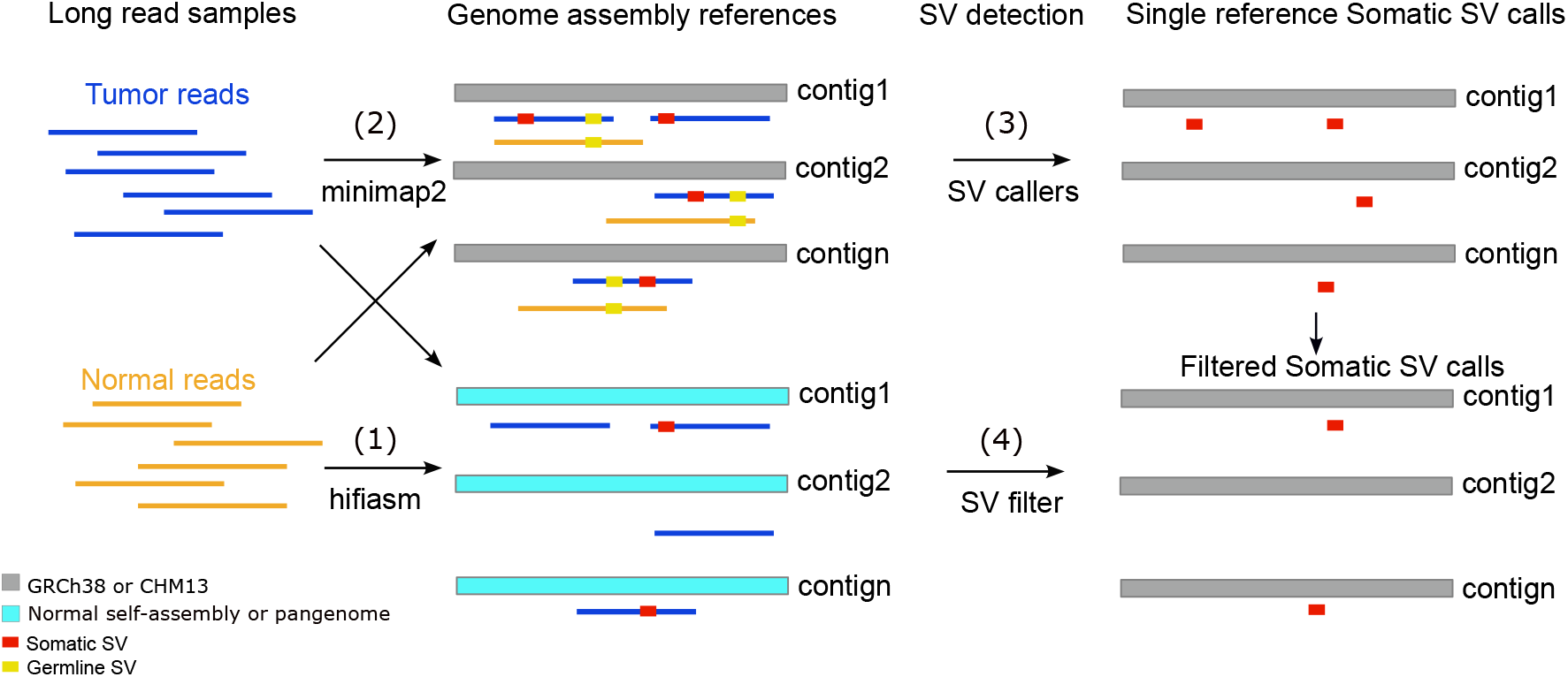
Overview of somatic SV calling and filtering workflow. Step (1): De novo assembly reads from the normal sample (if available). (2) Align both tumor and normal reads to GRCh38, self-assembly and/or pangenome. (3) Call candidate SVs from the GRCh38 alignment. (4) For each SV call, extract supporting reads and filter a read if its alignment to the self-assembly and/or pangenome does not contain the SV.

### Producing self-assembly

For a pair of matched tumor and normal samples, the normal sample is de novo assembled with hifiasm v0.19.6 given PacBio High-Fidelity (HiFi) reads. This assembly is called self-assembly. A more recent hifiasm v0.24.0 is used for the assembly of Oxford Nanopore (ONT) simplex reads with option “--ont”.

### Aligning long read sequences

HiFi reads were aligned to GRCh38, T2T-CHM13v2, and self-assembly by minimap2 (v2.28) with option “-ax map-hifi -s50” and aligned to the CHM13 pangenome graph by minigraph (v0.21) with option “-cxlr”. Long read alignments are stored in PAF and GAF formats. ONT reads were aligned to the same set of references but with a different minimap2 preset “lr:hq”.

### Extracting candidate SVs

The *extract* subcommand of minisv takes unsorted long-read alignment as input and detects candidate SVs in each read based on indels indicated by alignment CIGAR and breakpoints from split alignment records. By default, it ignores an alignment if it meets one of following conditions: 1) low fraction of aligned bases (<0.7), short alignment length (<100bp), low alignment length on each end (<2000bp), and low alignment length in the middle (<50bp); 2) low mapping quality (<5) and low mapping quality on each end (<30); 3) excessive number of ≥50bp indels per read (≥3 per 10kb). Indels spanning two different pangenome segments are also ignored. These thresholds aim at filtering extreme cases and do not greatly affect final results.

Minisv classifies each candidate SV to deletion, insertion, inversion, duplication, translocation, or breakend (BND). Each candidate SV is further annotated by chromosome (or graph genome contig), start, end, strand, breakpoints offset on the reference and the read, mapping quality, distance to centromere, read name, and SV type. For extracting SV signatures from self-assemblies, we allowed the mapping quality of long read alignment to be zero.

### Integrating candidate SVs from multiple genomes

The *isec* subcommand of minisv combines candidate SVs extracted from alignment against multiple genomes or pangenomes, which in this article include GRCh38 (labeled as “l” for linear), T2T-CHM13v2 (“t” for telomere-to-telomere), CHM13 graph (“g” for graph) and self-assembly (labeled as “s”). Suppose a candidate SV is observed at position *p* of a read in its alignment to GRCh38. The *isec* subcommand retains the SV on the read coordinate if the SV is observed within the 2kb window around position *p* on the same read in its alignment against another genome or pangenome; *isec* drops the candidate SV otherwise.

### Calling SVs supported by multiple reads

So far one unique candidate SV at a genomic position may be extracted multiple times. To combine reads supporting the same SV, the *merge* subcommand takes coordinate-sorted candidate SVs as input and merges two SVs if the two end points of one SV is within 100bp windows around the two end points of the other SV. If the SV is an insertion, the subcommand additionally requires the insertion length of the two SVs are within 5% of each other. On real data, somatic SVs are sparse and SVs of the same type are rarely close to each other. The SV calling results are insensitive to the thresholds.

Given a tumor-normal sample pair, the *merge* subcommand jointly considers both tumor and normal reads and counts the number of tumor and normal reads supporting each SV. An SV only supported by tumor reads but no normal reads is considered as a somatic SV. SVs supported by one read are discarded.

Because a pangenome encodes most common germline SVs, a read that contains an SV already present in the pangenome will be aligned without SVs. As a result, an SV called from pangenome alignment is either a rare SV or a somatic SV. Minisv leverages this observation to call mosaic SVs from a single sample, though it is unable to distinguish somatic SVs from rare ones.

### Simulating mosaic SVs

Tumor HiFi reads are downsampled with samtools and mixed with normal HiFi reads to a ratio of 1:10 on average across the genome. The tumor-to-normal ratio of a particular SV may deviate from the average due to statistical fluctuation. The simulated mix is assembled to produce the self-assembly.

### Generating somatic SV calls with other SV callers

Other callers were applied to the alignment against GRCh38 to call somatic SVs. Severus (v1.6) requires phased alignment as input. Following its documentation, we called SNPs and short INDELs in normal samples with Clair3 using option “--enable_phasing --longphase_for_phasing”. We used pre-trained model “hifi_revio” for HiFi and “r1041_e82_400bps_sup_v430” for ONT. We then applied “whatshap haplotag” to generate phased alignment. We ran Severus in the VNTR-aware mode, which generally improved SV accuracy. We used the same phased alignment as input to SAVANA (v1.3.6). Nanomonsv (v0.8.0) and Sniffles2 (v2.6.3) do not use phasing. We ran Sniffles2 under option “--tandem-repeats” to generates two intermediate “snf” files for tumor and normal samples, respectively. We then produced a two-sample VCF with Sniffles2 and selected tumor-only SV calls with “minisv.js snfpair”.

### Generating mosaic SV calls with other SV callers

Sniffles2 support mosaic SV calling with option --mosaic. Severus does not natively support mosaic SV calling. We instead ran Severus in the single-sample mode with and without panel of normal (PON) filtering and only retained SV calls with variant allele fraction below 25%.

### Filtering SV calls by other callers using self-assembly

While minisv can integrate alignment against self-assembly to filter out SVs not observed in self-assembly, other callers do not have this functionality built in. To improve other SV callers, we asked other SV callers to output the names of reads supporting each SV call, aligned the reads to the self-assembly and extracted candidate SVs with minisv. We filtered out a read (and thus reduced the supporting read count by one) if the read does not contain an SV in self-assembly alignment that does not match the original SV call.

Minisv considers an SV called from self-assembly (called “self” in brief) matching the original SV call by an SV caller (called “original”), if one of the following conditions is met: 1) both original and self SVs involve two chromosomes or contigs (i.e., both are translocations); 2) the original SV is longer than 100kb and the self SV involves two contigs, which could be caused by an assembly gap; or 3) both original and self SVs are not translocations and they are similar in length (differences between the two SV lengths are less or equal than 1kb, or SV length in one reference larger or equal than 60% of SV length in the other reference).

### Evaluating SV calls

The *eval* subcommand of minisv evaluates the accuracy of an SV call set against a truth SV set. It considers two SVs to be *equivalent* if 1) the end points of one SV are within 500bp windows around the end points of the other SV and 2) in case of INDELs, the difference in length is below 1kb or 40% of the longer SV. These rules are similar to the *merge* subcommand, but the thresholds are more relaxed because different SV callers may output SVs at slightly different locations. We manually checked alternative thresholds and found the default can reliably separate correct SV calls from wrong ones. Under this equivalence definition, a truth SV without equivalent test SVs is a false negative (FN), while a test SV without equivalent truth SV is a false positive (FP). The *eval* subcommand also optionally takes a list of confident regions, which in this article exclude centromeric regions, acrocentric short arms and pseudo-autosomal regions on chromosome X and Y. SV calls overlapping with these regions are ignored.

### Constructing ensemble call sets

The *union* subcommand constructs an ensemble call set from multiple input SV call sets. Conceptually, each SV call is considered as a node in a graph and two SVs are connected by an edge if they are equivalent according to the rules above. The subcommand performs single-linkage clustering to group SV calls from different callers. It then checks and reports whether all SV calls in a cluster are equivalent to each other. As somatic SVs are sparse in the genome, 99% of clusters are cliques – all SVs in a cluster are equivalent to each other but are not equivalent to SVs outside the cluster. This suggests single-linkage clustering works well in practice.

### Code and Data Availability

The JavaScript version of minisv is available at https://github.com/lh3/minisv, with additional functionality implemented in https://github.com/qinqian/minisv.py. The other SV callers, aligners, converter, and assembler software along with the dependent data of the VNTR BED file and minisv *eval* genome region BED file included in this study are listed in the table (Supplementary Table 1). The source codes to process, evaluate, and benchmark different SV callers are available at https://github.com/qinqian/pangenome_sv_benchmarking. The genome references used in the evaluation are listed in the summary table (Supplementary Table 1).

Both of HCC1395 and COLO829 cell lines along with their paired normal cell lines HiFi sequencing reads were obtained from the PacBio Cloud website. The rest of the cancer cell lines and paired lymphoblast normal cell lines PacBio HiFi and ONT R10 sequencing data were obtained from NCBI SRA database (SRP494767(10)). The paired normal sample PacBio HiFi sequencing data from the human pangenome reference project are available from the NCBI SRA database (SRP305758(16)) and Amazon Cloud. All the involved data are listed in the summary table (Table 1), we downloaded these SRA datasets using fastq-dump. All the SV calls and de novo assemblies are deposited to Zenodo (DOI: 10.5281/zenodo.17317401) https://zenodo.org/records/17317401.

**Table 1.**
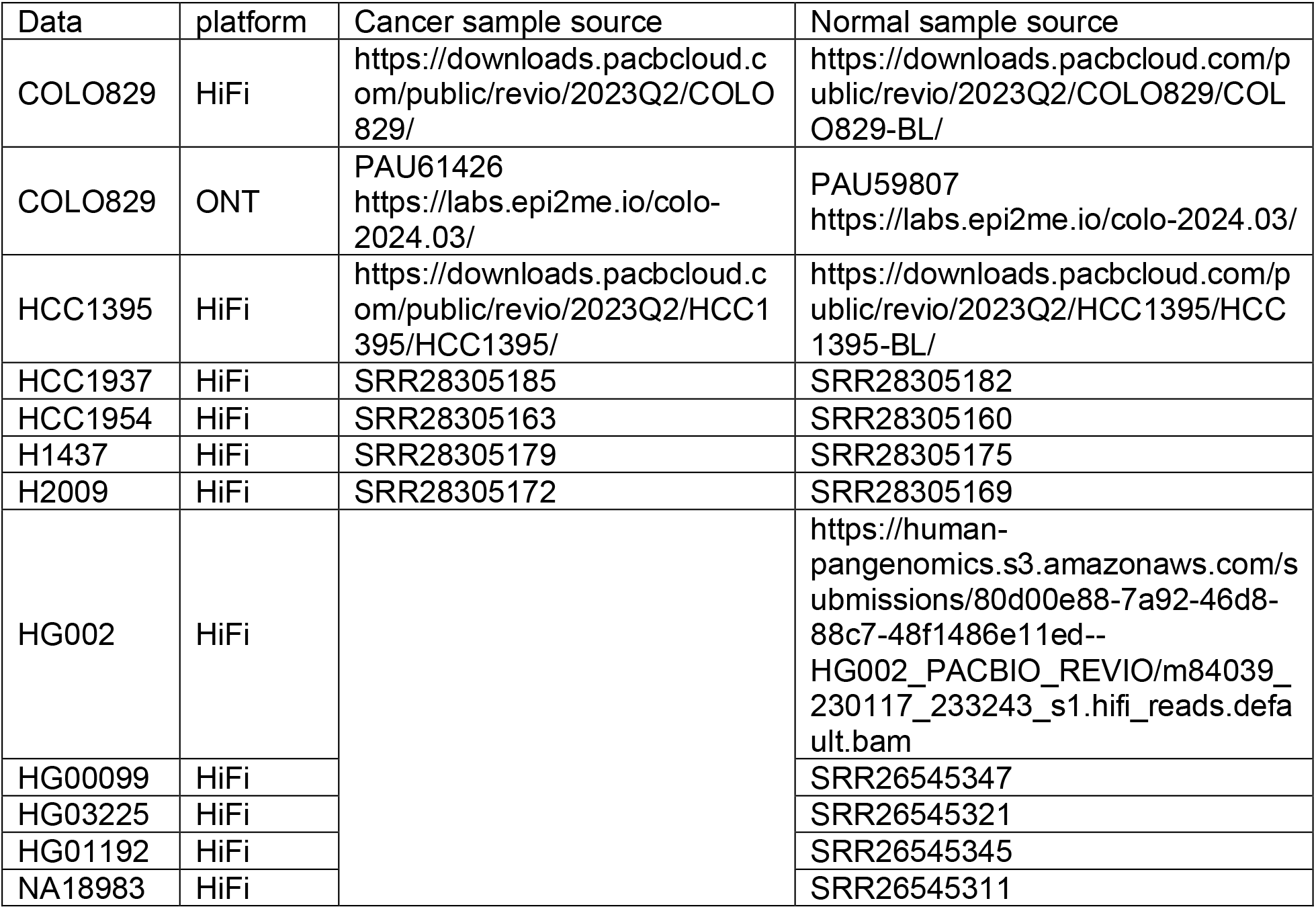
Summary of PacBio HiFi and Nanopore ONT sequencing data.

## Results

### Measuring false positive SVs with germline samples

While large SVs including ≥100kb indels and translocations are common in cancer, they are rare in germline. As a result, most large germline SVs are false positives. We can use this observation to measure the specificity of SV callers. We applied minisv and several other mainstream SV callers to the HiFi data of HG002 and four other lymphoblastoid samples from the human pangenome reference consortium (HPRC) (Table 1). GRCh38 was used as the reference genome. For minisv, we additionally combined the alignments against multiple genomes.

We observed that jointly considering GRCh38 (labeled by “l”) and CHM13v2 (labeled by “t”) resulted in 25-30 large SVs (Figure 2A). Adding graph genome (labeled by “g”) reduces the number of large SVs below 5. Most of the filtered SVs are translocations, which are potentially caused by smaller germline SVs not present in GRCh38. Further integrating self-assembly (labeled by “s”) filtered removes almost all large SVs except two cases in HG01192 and NA18983. Both SVs occurred at the IGL locus and are likely real somatic VDJ deletions. In comparison, Severus and Sniffles2 called 10-60 large SVs (Figure 2B). Almost of all of them are likely false positive calls given that large SVs are rare in germline and these SVs are not observed in the self-assembly.

**Figure 2.**
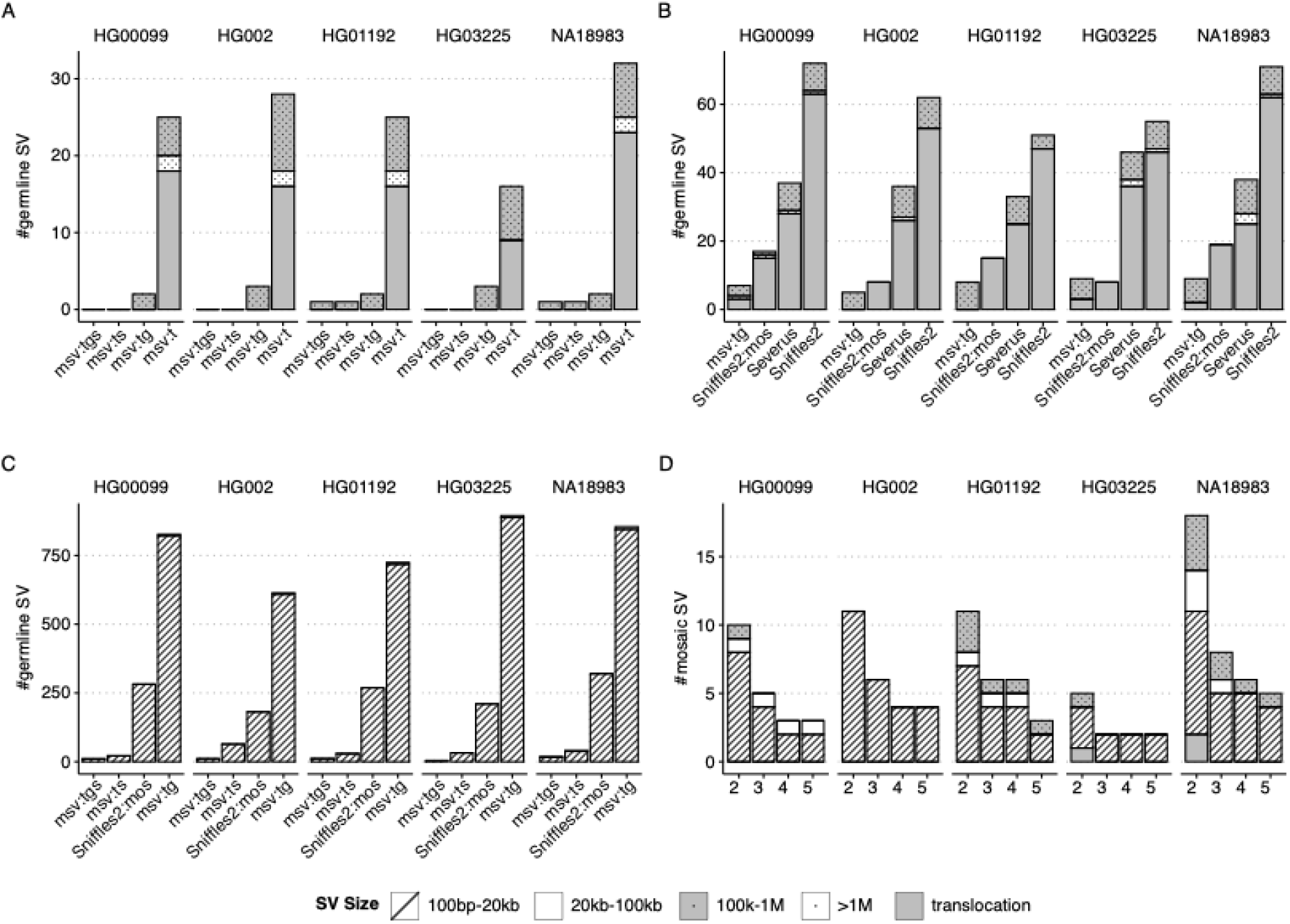
Germline and mosaic SV calling from normal cell lines. **A.** Long germline minisv calls (INDELs over 100kb in size or translocations) from different alignment combinations: “msv:t” for SVs called from GRCh38 and T2T-CHM13 alignment; “msv:tg” from alignment against GRCh38, T2T-CHM13 and pangenome graph; “msv:ts” from alignment against GRCh38, T2T-CHM13 and self-assembly; “msv:tgs” from alignment against all genomes/pangenome mentioned above. **B.** Long germline SV calls by different SV variant callers, Sniffles2:mos for mosaic mode using Sniffles2. **C.** Potential mosaic SV calls by different callers. **D.** Potential mosaic SV calls by msv:tgs. A zoom-in.

The GRCh38-CHM13-graph combination (labeled by “tg”) called 700-900 small SVs (100bp– 100kb) supported by ≥2 reads (Figure 2C). Most of these are rare SVs missing from the pangenome. Adding the self-assembly (labeled by “tgs”) drastically reduced the number to below 20, or lower if we require more supporting reads (Figure 2D). This demonstrates the power of self-assembly in mosaic SV calling.

In the mosaic mode, Sniffles2 called 190 candidate somatic SVs supported in HG002 (Figure 2C). Most of the SV-supporting reads are aligned to the HG002 diploid assembly without SVs. If we remove these reads, the number of somatic SV calls are reduced by 98% to 11. This suggests almost all of mosaic SVs called by Sniffles2 are either germline SVs or false SVs caused by germline variants relative to GRCh38.

### Evaluating somatic SV calling with COLO829 tumor-normal sample pairs

We took a previously established COLO829 callset (17) as the ground truth to test and quantify the effect of self-assembly on somatic SV calling. This callset consists of 58 SVs, including 38 indels ≥100bp, 7 foldback inversions and 13 translocations.

We called somatic SVs from the COLO829 tumor-normal pair with minisv in the “tg” mode (i.e. combining GRCh38, CHM13 and the CHM13 graph), nanomonsv, SAVANA and Severus. We filtered an SV call if the SV-supporting reads are aligned without SVs against the self-assembly (Methods). This filter barely affects true positive somatic SV calls, but it greatly reduces the number of false positive calls for all callers (Fig. 3). Between SV callers, minisv, nanomonsv, SAVANA and Severus have broadly similar accuracy. Note that we did not include Sniffles2 in this figure because it generated many false positive SVs without the self-assembly filter (Fig. 5).

**Figure 3.**
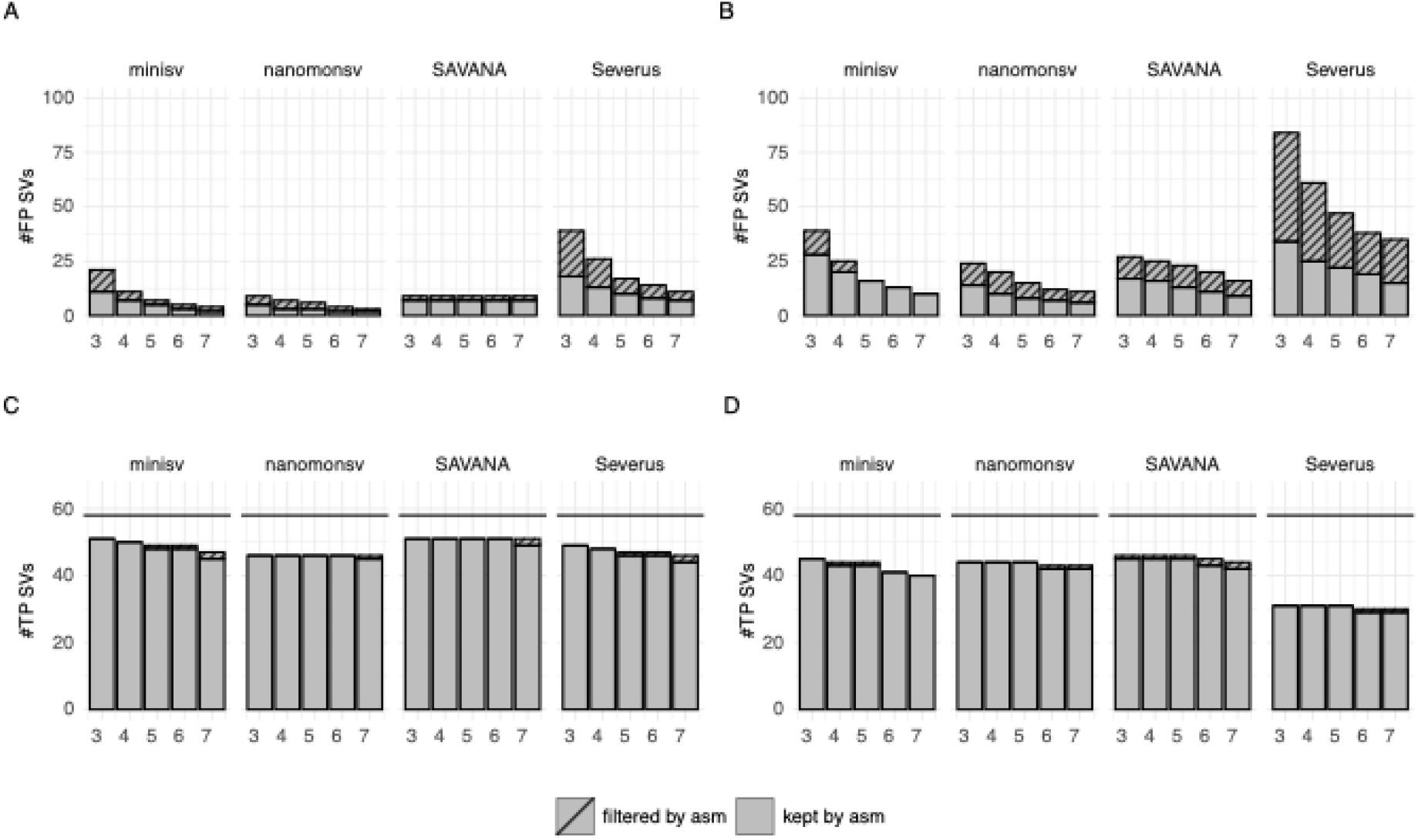
Somatic SV calling on matched COLO829 tumor-normal cell lines. **A.** Number of false positive (FP) and true positive (TP) SV calls from PacBio HiFi data at different cutoffs on supporting reads. The solid bars correspond to SV calls still observed in the alignment against the COLO829BL (derived from blood) assembly, while the shaded bars correspond to calls filtered by the assembly. The horizontal line gives the number of SV calls from Valle-Inclan et al., which are taken as ground truth. **B.** Number of FP and TP calls from ONT reads.

We noticed that no callers could call all SVs in the truth set and that even with the self-assembly filter, there are still a small number of SV calls not found in the truth set. We reasoned that the COLO829 cancer cell line may be still evolving across different passages of cell growth (18,19) and different data centers may have generated sequencing data with different DNA materials. This may also explain the slightly higher number of extra and missing ONT calls after the self-assembly filter (Fig. 3B).

### Emulating and evaluating mosaic SV calling with COLO829 data

To emulate a dataset with known mosaic SVs, we downsampled the COLO829 tumor reads and mixed them with the normal reads *in silico* such that the normal-to-tumor ratio is 10:1 on average. Given approximately 60-fold tumor and normal read coverage in our data, the tumor read coverage is six across the genome. As each somatic SV only occurs to one haplotype in tumor cells, the number of reads supporting the SV is only three on average. Although some SVs may be supported by more reads due to aneuploidy and focal amplifications, we still expect SV callers to miss many somatic SVs in the in-silico mix.

We called mosaic SVs with minisv in the “tg” mode, Sniffles2 in the mosaic mode, and Severus with post hoc allele fraction filter (Methods). We required each SV call to be supported by ≥2 reads. Using alignment against GRCh38 only, all three tools called hundreds of supposedly somatic SVs (Fig. 4). Most of these SVs can be found in the normal sample – they are germline SVs and are thus false positive somatic calls. The self-assembly filter reduces the number of such false positives by more than an order of magnitude while retaining the great majority of true positive calls. The several true somatic SVs mistakenly filtered by the self-assembly tend to reside in regions with focal amplification in COLO829. They are supported by enough tumor reads to be assembled into contigs. The self-assembly filter may affect sensitivity for mosaic SVs of high allele fraction.

**Figure 4.**
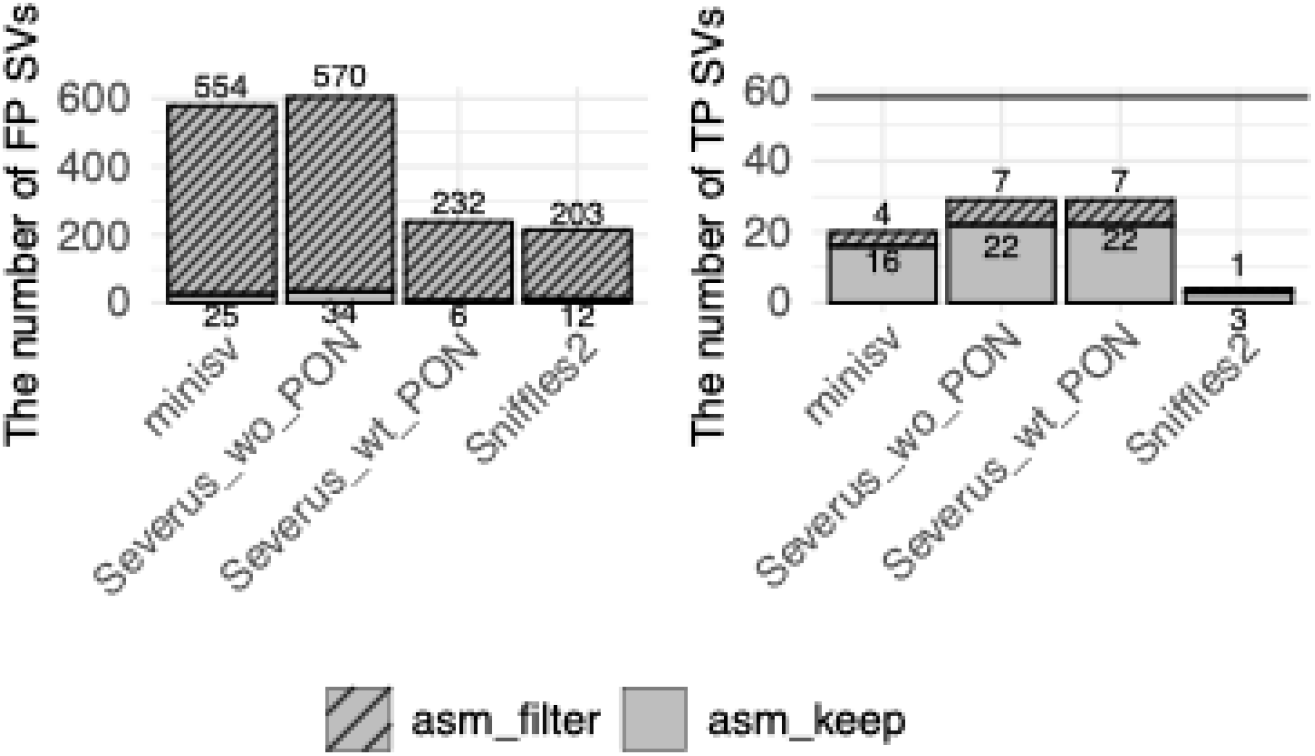
Mosaic SV calling from in-silico mix of normal-tumor data. COLO829 tumor HiFi reads were computationally downsampled and mixed with all COLO829BL normal reads to 10:1 normal-to-tumor ratio. The in-silico mix was assembled and used for assembly-based filtering.

Between SV callers, Severus with our filtering has the best sensitivity. Sniffles2 in the mosaic mode misses most somatic SVs identified by other tools.

### Investigating somatic SV calling for additional tumor-normal pairs

We ran all SV callers on five other tumor-normal paired cell lines including HCC1395, HCC1937, HCC1954, H1437 and H2009. As there are no curated ground truth for these cell lines, we could not compute true positives and false positives, but we could see the self-assembly filter continued to play a role in reducing somatic SV calls especially for SVs supported by few reads (Fig. 5). These filtered SVs are likely false positives. The number of SVs called by each tool, except Sniffles2, is broadly comparable after the self-assembly filter.

**Figure 5.**
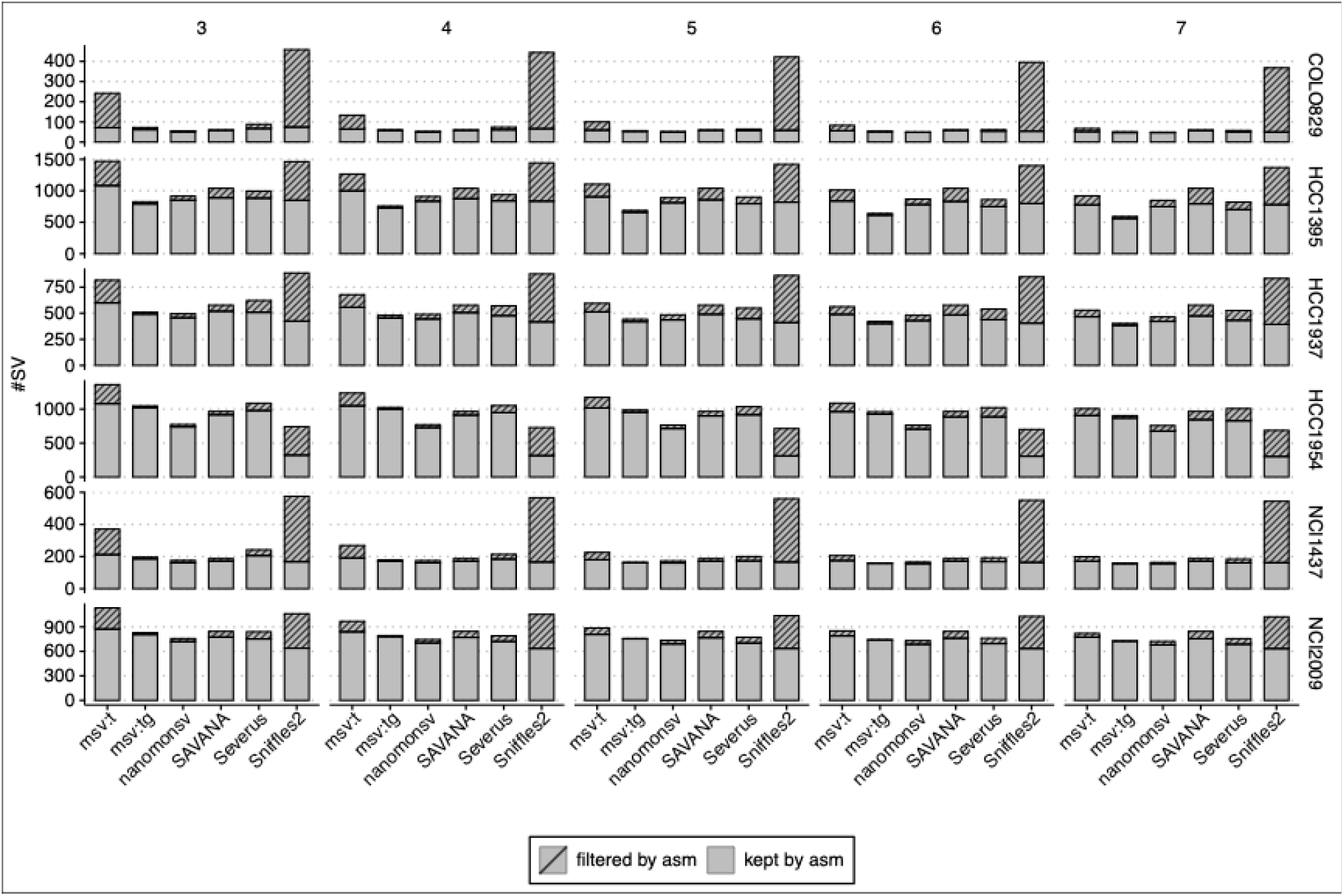
Effect of assembly-based filtering on SV calling on six tumor-normal pairs. The number on the top indicates the number of supporting reads required to call an SV.

We ran minisv in two modes, “lt” and “ltg”, with the former considering the linear reference genomes only and the latter additionally considering the pangenome alignment. Using a simple algorithm, the “lt” mode of minisv did not compete with other sophisticated callers in accuracy. Nevertheless, including the pangenome, the “ltg” mode produced fewer false positives than other callers. This demonstrates the effectiveness of pangenome. Minisv “ltg” missed more somatic SVs for HCC1395 because many somatic SVs in this sample appeared to fall in tandem repeats which often overlapped with germline SVs in the pangenome and thus got filtered.

Minisv, nanomonsv, SAVANA and Severus generally agree each other with most somatic SVs identified by all four callers (Fig. 6). The great majority of SVs called by all four tools are also supported by self-assemblies and are likely to be accurate. On the contrary, SVs called by one or two callers are less reliable.

**Figure 6.**
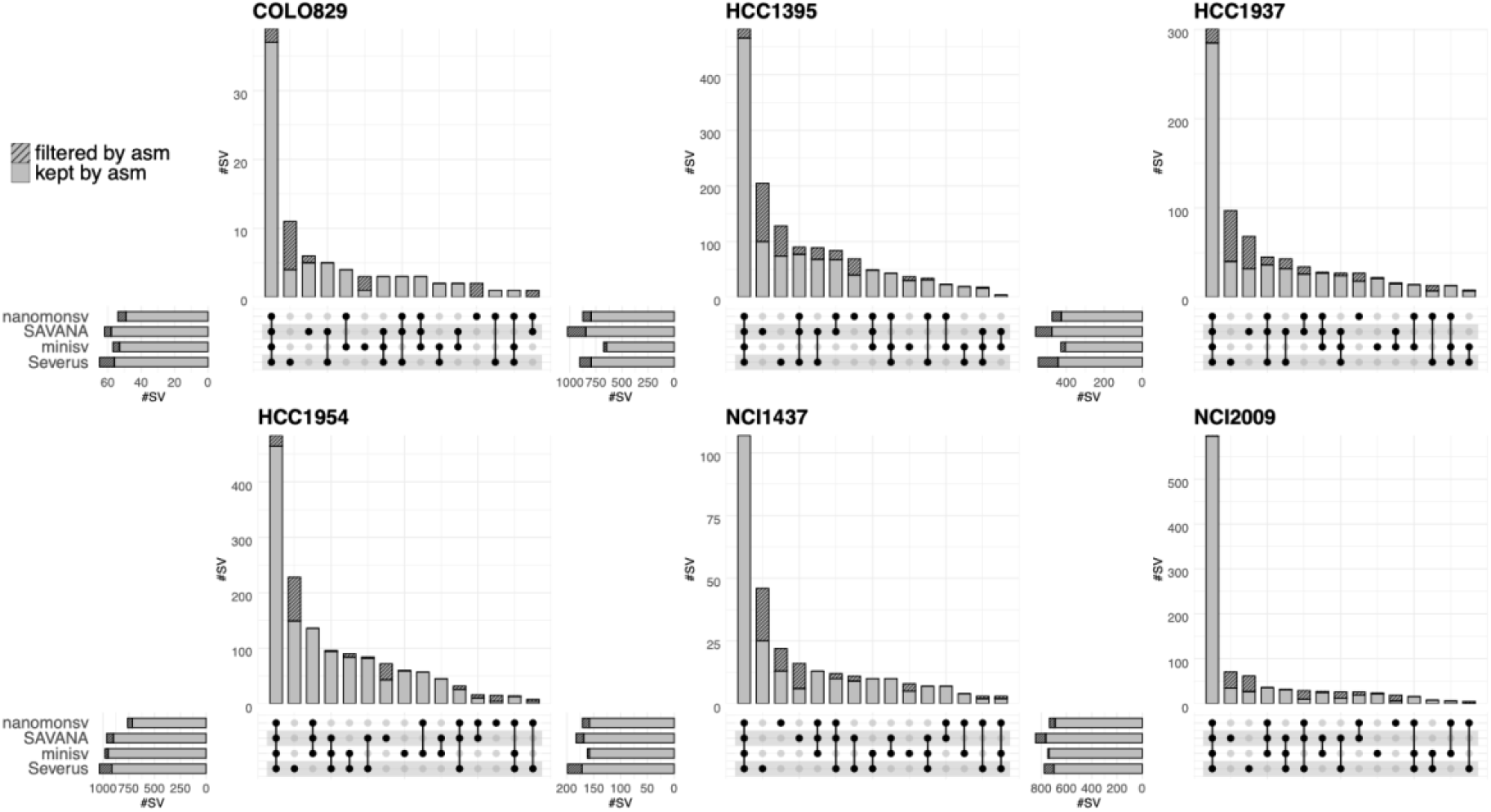
Intersection of somatic SVs called by four callers from six tumor-normal pairs.

## Discussions

Implemented in minisv, our method is unique in its use of alignment against multiple genomes including pangenome and the assembly of normal samples. It greatly reduces alignment artifacts caused by germline SVs relative to the reference genome, and thus improves the specificity of somatic or mosaic SV calls at little loss in sensitivity. Minisv works as a standalone tool or can be used to filter false positive SVs called by other callers. It also comes with features rarely found in other toolkits, such as SV evaluation and the construction of union callsets for translocations and foldbacks.

Applying our method to multiple datasets, we found that SV callers specialized for tumor-normal pairs, which include nanomonsv, Severus, SAVANA and minisv, all work reasonably well in that the majority of calls are shared by all callers. Self-assembly-based filtering improves the accuracy but not dramatically. Calling mosaic SVs of low allele fraction is much more challenging. All current methods would report many germline or artifactual SVs as mosaic. The self-assembly filter is particularly powerful in this setting and is highly recommended.

A potential concern with the self-assembly filter is that somatic SVs of high allele fraction may be represented in the assembly and may be mistakenly filtered with our method. A better solution is to keep a candidate somatic SV call if it only occurs to one haplotype. Taking the advantage of haplotype phasing will also improve mosaic SV calling in normal tissues. Another weakness of our current implementation is that we need to align all reads against multiple genomes or pangenomes. This will increase the total runtime but can be alleviated by only aligning reads containing candidate SV calls. We are developing an integrated long-read variant caller that will address these technical issues and supposedly achieve higher accuracy and performance.

## Authors’ Disclosures

No disclosures were reported.

## Authors’ contributions

H. Li: Conceptualization, supervision, methodology, software, data analysis, data acquisition, manuscript writing, funding acquisition.

Q. Qin: Methodology, software, data analysis, data curation, data acquisition, manuscript writing J. Heinz: Software, data analysis.

## Acknowledgments

The authors are grateful to Ying Zhou, Megan Le, Yan Gao for the feedback, and to Chengzhong Zhang for the discussions. This work was supported by grants from the US National Institutes of Health (U24CA294203, R01HG010040 and U01HG013748 to H.L.).

## Supplementary Tables and Figures

Supplementary Table 1. Summary of software source codes and reference genomes.

## Notes

### Competing Interest Statement

The authors have declared no competing interest.

## References

1. Quinlan AR, Hall IM. Characterizing complex structural variation in germline and somatic genomes. Trends in Genetics. 2012. page 43–53.

2. Raffaele Cosenza M, Rodriguez-Martin B, Korbel JO. Structural Variation in Cancer: Role, Prevalence, and Mechanisms SNV: single-nucleotide variant (here used largely for somatic variants). Annual Review of Genomics and Human Genetics Downloaded from http://www.annualreviews.org Guest [Internet]. 2024;17:44. Available from: 10.1146/annurevgenom-120121-

3. Hanahan D, Weinberg RA. Hallmarks of cancer: the next generation. Cell [Internet]. Elsevier Inc.; 2011;144:646–74. Available from: http://www.ncbi.nlm.nih.gov/pubmed/21376230

4. Campbell PJ, Getz G, Korbel JO, Stuart JM, Jennings JL, Stein LD, et al. Pan-cancer analysis of whole genomes. Nature. Nature Research; 2020;578:82–93.

5. Li Y, Roberts ND, Wala JA, Shapira O, Schumacher SE, Kumar K, et al. Patterns of somatic structural variation in human cancer genomes. Nature. Nature Research; 2020;578:112–21.

6. Dohm JC, Lottaz C, Borodina T, Himmelbauer H. Substantial biases in ultra-short read data sets from high-throughput DNA sequencing. Nucleic Acids Res. 2008;36.

7. Benjamini Y, Speed TP. Summarizing and correcting the GC content bias in high-throughput sequencing. Nucleic Acids Res. 2012;40.

8. Marx V. Method of the year: long-read sequencing. Nat Methods. 2023;6– 11.

9. Chaisson MJP, Huddleston J, Dennis MY, Sudmant PH, Malig M, Hormozdiari F, et al. Resolving the complexity of the human genome using single-molecule sequencing. Nature. Nature Publishing Group; 2015;517:608–11.

10. Keskus A, Bryant A, Ahmad T, Yoo B, Aganezov S, Goretsky A, et al. Severus: accurate detection and characterization of somatic structural variation in tumor genomes using long reads [Internet]. 2024. Available from: http://medrxiv.org/lookup/doi/10.1101/2024.03.22.24304756

11. Elrick H, Sauer CM, Espejo Valle-Inclan J, Trevers K, Tanguy M, Zumalave S, et al. SAVANA: reliable analysis of somatic structural variants and copy number aberrations in clinical samples using long-read sequencing [Internet]. 2024. Available from: http://biorxiv.org/lookup/doi/10.1101/2024.07.25.604944

12. Lin J, Wang S, Audano PA, Meng D, Flores JI, Kosters W, et al. SVision: a deep learning approach to resolve complex structural variants. Nat Methods. Nature Research; 2022;19:1230–3.

13. Smolka M, Paulin LF, Grochowski CM, Horner DW, Mahmoud M, Behera S, et al. Detection of mosaic and population-level structural variants with Sniffles2. Nat Biotechnol. Nature Research; 2024;

14. Shiraishi Y, Koya J, Chiba K, Okada A, Arai Y, Saito Y, et al. Precise characterization of somatic complex structural variations from tumor/control paired long-read sequencing data with nanomonsv. Nucleic Acids Res. Oxford University Press; 2023;51:E74.

15. Ghandi M, Huang FW, Jané-Valbuena J, Kryukov G V., Lo CC, McDonald ER, et al. Next-generation characterization of the Cancer Cell Line Encyclopedia. Nature. Nature Research; 2019;569:503–8.

16. Liao WW, Asri M, Ebler J, Doerr D, Haukness M, Hickey G, et al. A draft human pangenome reference. Nature. Nature Research; 2023;617:312–24.

17. Espejo Valle-Inclan J, Besselink NJM, de Bruijn E, Cameron DL, Ebler J, Kutzera J, et al. A multi-platform reference for somatic structural variation detection. Cell Genomics. Cell Press; 2022;2.

18. Paulin LF, Fan J, O’Neill K, Pleasance E, Porter VL, Jones SJM, et al. The benefit of a complete reference genome for cancer structural variant analysis [Internet]. 2024. Available from: http://medrxiv.org/lookup/doi/10.1101/2024.03.15.24304369

19. Velazquez-Villarreal EI, Maheshwari S, Sorenson J, Fiddes IT, Kumar V, Yin Y, et al. Single-cell sequencing of genomic DNA resolves sub-clonal heterogeneity in a melanoma cell line. Commun Biol. Nature Research; 2020;3.

